# Tracking the progression of Alzheimer’s disease with peripheral blood monocytes

**DOI:** 10.1101/2023.02.28.530459

**Authors:** Viktoriia Bavykina, Mariano Avino, Mohammed Amir Husain, Adrien Zimmer, Hugo Parent-Roberge, Abdelouahed Khalil, Marie A. Brunet, Tamas Fülöp, Benoit Laurent

**Author notes:** Corresponding authors (B. Laurent), (T. Fülöp).

## Abstract

**Background:** Alzheimer’s disease (AD) is the most common form of dementia with the symptoms gradually worsening over the years. However, the driving pathological processes occur well before the appearance of symptoms. AD patients display signs of systemic inflammation, suggesting that it could precede the well-established AD hallmarks. We recently showed that the innate immune response in the form of monocyte activation is detectable at the pre-clinical stage.

**Objectives:** Our goal here is to characterize changes of gene expression in peripheral blood monocytes from patients at different stages of AD progression and validate potential biomarkers for a better prognosis and diagnosis of AD clinical spectrum.

**Results:** We performed a whole transcriptome analysis on monocytes purified from healthy subjects, Mild Cognitive Impairment (MCI) and AD patients, and established the list of genes differentially expressed in monocytes during the disease evolution. We observed that, in the top 500 genes differentially expressed, a majority of these genes were upregulated (65%) during AD progression. These genes are mainly involved in chemokine/cytokine-mediated signaling pathways. We further confirmed several biomarkers by quantitative PCR and immunoblotting and showed that they are often deregulated at pre-clinical stages of the disease (MCI stage), supporting the hyperactivation of monocytes in MCI patients.

**Perspectives:** Our findings provide evidence that the pre-clinical stage of AD can be detected in monocytes using a specific set of biomarkers, highlighting the importance to study the early innate immune response in AD. Our results open the possibility to use these biomarkers with different diagnostic methodologies to better predict and efficiently treat AD.

## Introduction

Alzheimer’s disease (AD) is an irreversible neurodegenerative disease manifesting ultimately as clinical dementia. AD is initiated decades before its clinical diagnosis, suggesting that the driving pathological processes occur well before the manifestation of symptoms. Before the appearance of the well-established AD hallmarks (i.e., deposition of Aβ plaques and neurofibrillary tangles), AD condition displays signs of a systemic inflammation ^1-3^. The innate immune system assures the first line of defence against internal and external attacks (e.g. infections, vascular and metabolic insults), but the accumulation of threats fosters the permanent activation of innate immune cells hence contributing to low but significant secretion of pro-inflammatory mediators ^4^. In the brain, this progressive neuroinflammation causes irreversible damages such as the blood-brain barrier breakdown, allowing inflammatory mediators to reach the periphery and trigger the systemic inflammatory response observed in AD patients ^5, 6^. Since AD pathogenesis exhibits early signs of a systemic inflammation, the peripheral immune response should change with the progression of the disease. We showed that peripheral monocytes are activated at early stages of the disease, indicating that these cells could be good candidates for the discovery of biomarkers assessing AD progression ^7^. Here we report our investigation on the gene expression profile of peripheral monocytes from patients at different stages of the disease to identify biomarkers that could help us to track the evolution of the disease.

## Methods

### Participants

Mild Cognitive Impairment (MCI; n=9) and mild AD (AD; n=10) patients were recruited from the registry of the memory clinic of the Sherbrooke Geriatric University Institute. Healthy subjects (H; n=9) were recruited from our healthy individual database at the Research Center on Aging at Sherbrooke and by advertisement. All subjects gave written informed consent. The project was approved by the Institutional Review Board of the CIUSSS de l’Estrie-CHUS (Project #2019-2877 – ADAUDACE - Fülöp). Details concerning the characteristics of each subject group have been previously described ^7^. Demographic data, cognitive test results, hematological and biochemical values for each subject group were listed in **Table 1**.

**Table 1.** Clinical data of patients. MMSE: Mini-Mental State Examination; MoCA: Montreal Cognitive Assessment; ALT: alanine aminotransferase; AST: aspartate aminotransferase; GFR: glomerular filtration rate; WBC: white blood cells; TSH: thyroid stimulating hormone.

| Parameters |  | Healthy subjects<br>(H) (n=9) |  | MCI patients<br>(MCI) (n=9) |  | AD patients<br>(AD) (n=10) |  | p-values<br>Tukey HSD test |  |  |
| --- | --- | --- | --- | --- | --- | --- | --- | --- | --- | --- |
| Type | Units | mean | SD | mean | SD | mean | SD | MCI vs H | AD vs H | MCI vs AD |
| Age | years | 71.8 | 5.8 | 72.4 | 6.1 | 76.3 | 7.7 | 0,9993 | 0,3323 | 0,35 |
| Gender | women (%) | 44 |  | 33 |  | 60 |  |  |  |  |
|  | men (%) | 56 |  | 66 |  | 40 |  |  |  |  |
| MMSE | score / 30 | 29.2 | 1.2 | 29 | 1 | 23.7 | 3.6 | 0,9958 | <0,0001 | 0,0001 |
| MoCA | score / 30 | 27.8 | 2 | 26.1 | 2 | 13.6 | 10.2 | 0,8335 | 0,0001 | 0,0006 |
| ALT | IU/L | 15 | 4.9 | 18.8 | 6.4 | 18.3 | 5.2 | 0,6321 | 0,6999 | 0,9844 |
| AST | IU/L | 20.7 | 3.8 | 20 | 2.7 | 21.7 | 2.8 | 0,9450 | 0,8645 | 0,5604 |
| Total bilirubin | mmol/L | 7.4 | 1.4 | 10.5 | 2.3 | 10.2 | 4.1 | 0,3759 | 0,4510 | 0,9732 |
| Total cholesterol | μmol/L | 4.2 | 0.6 | 4.7 | 0.5 | 5.2 | 0.9 | 0,6009 | 0,1341 | 0,4069 |
| Creatinine | μmol/L | 62.3 | 8.1 | 69.2 | 9.5 | 72.6 | 12 | 0,6600 | 0,3740 | 0,8527 |
| GFR |  | 81.9 | 4.2 | 87.3 | 1.2 | 77.8 | 8.3 | 0,4696 | 0,6113 | 0,0518 |
| Hemoglobin | g/L | 126.7 | 5.8 | 144.5 | 9.2 | 142.6 | 12.4 | 0,0939 | 0,1344 | 0,9411 |
| WBC | 10<sup>9</sup>/L | 5.3 | 1.1 | 5.3 | 1.5 | 6.3 | 1.1 | 0,9936 | 0,5049 | 0,4290 |
| Glucose | mmol/L | 4.6 | 0.2 | 5.7 | 1.9 | 5 | 0.5 | 0,4769 | 0,9226 | 0,5645 |
| TSH | μU/ml | 1.7 | 0.3 | 2.5 | 0.8 | 3.4 | 1.1 | 0,4163 | 0,0392 | 0,2145 |

### Samples collection and preparation

Monocytes were isolated from the whole blood of participants as previously described ^7^. Cell viability was measured at >95% using trypan blue exclusion assay. Total RNA was isolated from each sample using the Quick-RNA™ Miniprep Kit (Zymo Research). RNA-seq libraries were prepared with 10 ng of total RNA using NEBNext® single cell/low input RNA library preparation Kit (New England Biolabs). Single-end sequencing was performed on all the samples with a 75 bp sequencing depth on a NextSeq machine at the RNomics platform of the Université de Sherbrooke (Canada). The raw sequencing data are deposited at the Gene Expression Omnibus (GEO) under the subseries entry GSE216918.

### Biomarker validation

Biomarker candidates were validated by quantitative PCR (qPCR) and immunoblotting. For qPCR, cDNA were synthesized using the iScript Reverse Transcription Supermix (Bio-Rad) following manufacturer’s instructions. The qPCR reactions were performed with the PerfeCTa SYBR Green SuperMix (QuantaBio) as recommended by the manufacturer. Real-time PCR was performed on the Azur Cielo 3 system (Azure Biosystems). Sequences of primers used for qPCR are available upon request. For immunoblotting, protein samples were prepared and analyzed as previously described ^7^ using the following antibodies: anti-CCR3 (Abclonal A20253), anti-MMP8 (Abclonal A1963), anti-TMEM176B (Abclonal A16118), anti-β-actin (Sigma-Aldrich A5441). Pictures were taken with an iBright 1500 imaging system (Thermo Fisher).

## Results

To establish a list of blood-based biomarkers, we first performed a transcriptomic analysis to compare gene expression in monocytes from healthy elderly individuals (H), Mild Cognitive Impairment (MCI) and Alzheimer’s Disease (AD) patients (n=2 for each subject group). We analyzed each individual gene and their differential expression during the AD continuum to establish a list of the most differentially expressed genes (DEG) between the healthy and AD stages. We represented these DEG in a volcano plot to illustrate that some genes were upregulated up to 6 times (e.g., Ifit3, IL6, and CSF3 genes), while some DEG were downregulated up to 4 times (e.g., TRNS1) (**Figure 1A**). Hierarchical clustering of the top 500 DEG showed that the transcriptomic signature of monocytes from healthy individuals was different from that of monocytes from MCI and AD patients (**Figure 1B**). This clustering also indicated that monocytes of MCI and AD patients had a similar gene expression profile, suggesting that an AD-specific transcriptomic signature could be established in monocytes at AD pre-clinical stages. Among these DEG, the majority was upregulated during the disease continuum (n=311) while some genes exhibited a decrease in their expression (n=189) (**Figure 1B**). We finally performed a gene ontology (GO) analysis to identify in which specific pathways or functions those genes were involved. Using ShinyGO, a tool for in-depth analysis of gene sets ^8^, we showed that upregulated genes (n=311) were mainly involved in biological process of chemokine-mediated signaling pathways and defense response to virus while the 189 down-regulated genes were mostly involved in lipid antigen binding (**Figure 1C-D**). Analyses of target gene-transcription factor associations with the ENCODE Transcription Factor Targets Database, that catalogues target genes of transcription factors from transcription factor binding site profiles, also revealed that genes whose expression was increased in monocytes during the progression of AD were mainly regulated by the STAT2 transcription factor while down-regulated genes were the target of the IRF3 transcription factor (**Figure 1E-F**).

**Figure 1.**
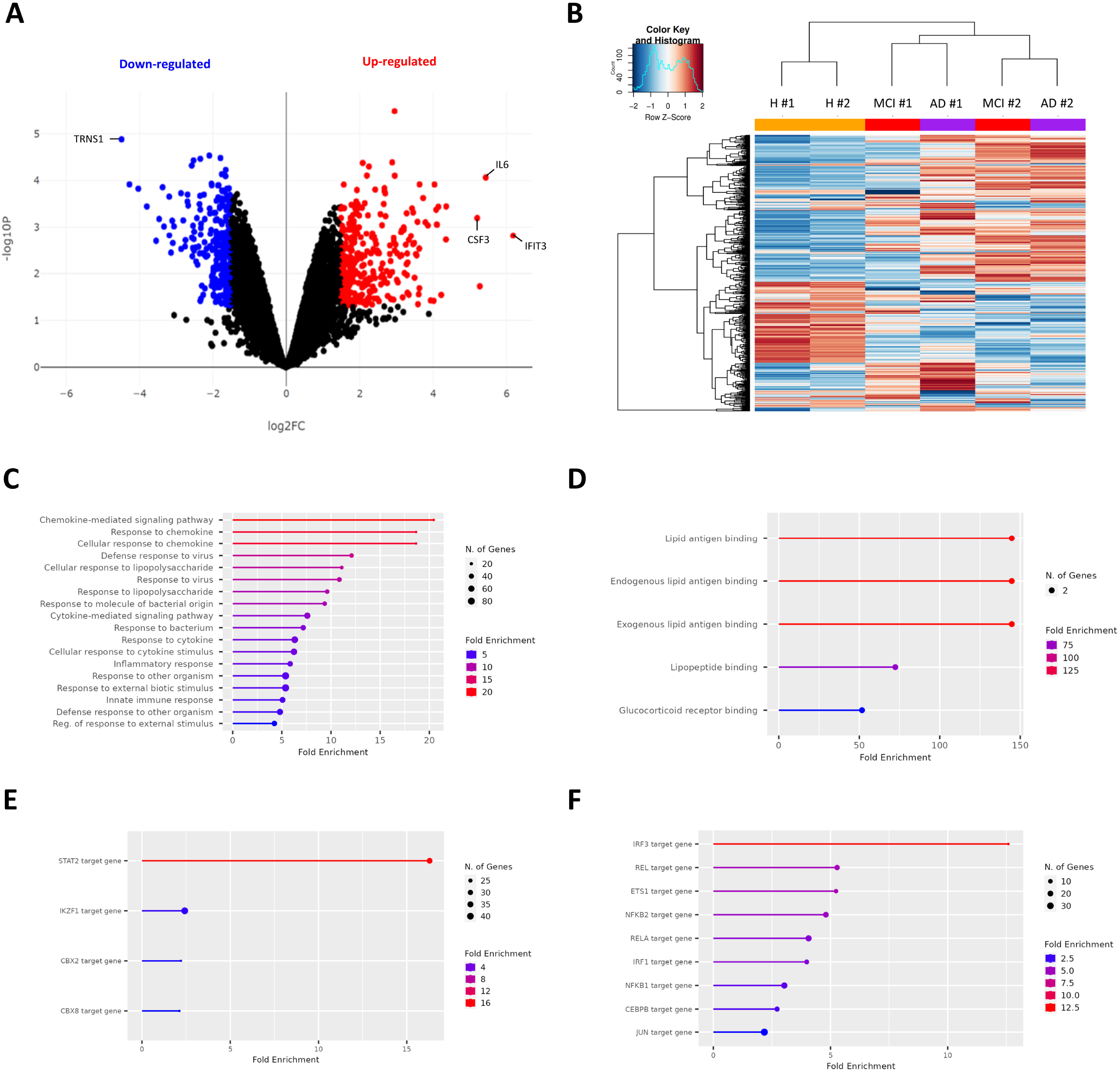
Changes in the transcriptome signature of monocytes during AD continuum. **(A)** Hierarchical clustering of the top 500 genes showing significant differential expression between healthy elderly individuals (H; orange), Mild Cognitive Impairment (MCI; red) and Alzheimer’s disease (AD; purple) patients (n=2 per subject group). Red and blue colors indicate relatively higher and lower expression, with genes independently scaled to a mean of zero. **(B)** Volcano plot of differentially expressed genes, mapping the upregulated genes (red) and downregulated genes (blue) at a 1% FDR using BioJupies tool ^17^. FC, fold change. **(C-D)** Gene Ontology analysis of upregulated **(C)** and downregulated **(D)** genes using the ShinyGO tool (version 0.76.1) ^8^. **(D-E)** Target gene-transcription factor associations by binding of transcription factor near transcription start site of upregulated **(D)** and downregulated **(E)** genes using the ENCODE Transcription Factor Targets Database.

To validate our transcriptomic results, we then performed quantitative PCR (qPCR) on more patients samples (n=9 per group). Among the DEG, we selected candidates with an overall good mRNA expression and/or with a biological relevance for the functions of interest e.g., chemokine-related genes. We picked a 10-gene panel of candidates that were downregulated (*LAMB3, USP18, NR4A1, IL1R2, CMPK2, S100B*) and up-regulated (*IFIT3, MMP8, LCN2, TMEM167B*) during the evolution of the disease. Our qPCR results confirmed the differentially expression of all genes in monocytes. We observed that some genes were progressively dysregulated during the progression of AS with, for instance, the expression of CMPK2 decreasing in average by 88% (109.8 ± 84.0 in healthy monocytes vs 13.6 ± 13.6 in AD monocytes) or the expression of TMEM167B increasing by 362% (3.7 ± 3.2 in healthy monocytes vs 13.4 ± 6.8 in AD monocytes) (**Figure 2A**). Interestingly, some candidate genes like IFIT3 exhibited an increase of gene expression specifically at the MCI stage (**Figure 2A**). Even we validated our 10-gene panel, it is important to note that there is a certain variability in gene expression among patients’ monocytes. As higher levels of mRNA have more chances to produce more proteins, we then investigated whether changes of mRNA level correlated with those of proteins. We examined by immunoblotting the expression of three upregulated genes (i.e., MMP8, CCR3, TMEM167B) in monocytes of healthy elderly individuals (H), MCI and AD patients, and showed that their protein levels were indeed significantly increased during the disease progression (**Figure 2B**). This increase was observed as early as the MCI stage, suggesting that an AD-specific signature is established at protein level in monocytes at AD pre-clinical stages.

**Figure 2.**
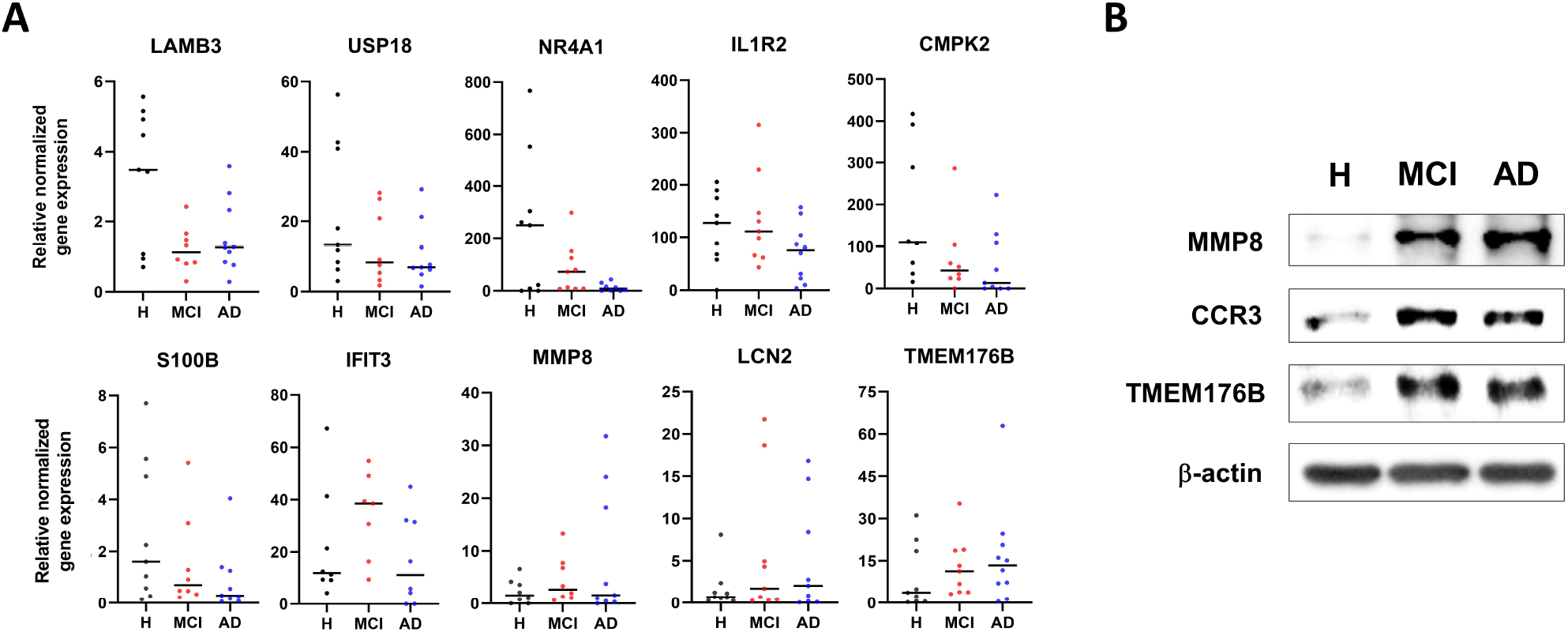
Validation of candidate biomarkers at mRNA and protein levels. **(A)** mRNA levels of IL1R2, USP18, CMPK2, NR4A1, S100B, LAMB3, MMP8, TMEM176B, LCN2, and IFIT3 were measured by quantitative PCR in monocytes of healthy elderly individuals (H; n=9), Mild Cognitive Impairment (MCI; n=9) and Alzheimer’s disease (AD; n=10) patients. mRNA levels were normalized to that of GAPDH. **(B)** Protein levels of MMP8, CCR3 and TMEM176B in monocytes of Healthy elderly control (H), Mild Cognitive Impairment (MCI) and Alzheimer’s disease (AD) patients were analyzed by immunoblot. Actin was used as a loading control.

## Discussion

Recent evidence highlighted neuroinflammation as a main hallmark of AD occurring decades before the appearance of clinical symptoms such as amyloid plaques and neurofibrillary tangles ^9^. It is now accepted that peripheral innate and adaptive immune cells infiltrate the AD brain as a result of the increased permeability of the blood-brain barrier. We recently showed that the innate immune response in the form of monocyte activation is detectable as early as the pre-clinical MCI stage ^7^. Our goal here was to assess the gene expression profile of peripheral monocytes purified from patients across the spectrum of AD, and potentially identify biomarkers to track the disease evolution. We established a list of genes differentially expressed in monocytes during AD evolution. Interestingly, we observed that 65% of these genes were upregulated (**Figure 1B**) and that their gene expression was progressively dysregulated as the disease progresses (**Figure 2A**). Changes in epigenetic mechanisms could potentially lead to a loss of chromatin structure, especially heterochromatin, during the progression of the disease ^10, 11^. Therefore, these changes could trigger alterations on the transcriptional level of specific gene subsets involved in monocyte activation and hence participate to the pathogenesis of AD ^12^. Epigenetic mechanisms play various roles in AD development so understanding the epigenetic contribution to monocyte activation and dementia pathogenesis may help to identify therapeutic targets for the inhibition of AD-associated trajectories ^13^.

We also determined that DEG were enriched in genes target for the STAT2 and IRF3 transcription factors (**Figure 1E-F**). Interestingly, the JAK/STAT signaling pathway is known to promote neuroinflammation in AD by initiating innate immunity and orchestrating adaptive immune mechanisms ^14^. Overexpression or increase activity of STAT2 might trigger a neuroinflammatory response during AD progression by upregulating a specific subset of genes. In another hand, the cGAMP-STING-IRF3 pathway induces the expression of TREM2, whose expression decreases Aβ deposition while improves AD cognitive impairment ^15^. Decrease of IRF3 expression and hence its downstream target genes might therefore increase Aβ deposition during AD progression and participate to the physiopathology. It will be important to investigate the status of these transcription factors in monocytes as the disease progresses.

We also showed that changes of mRNA expression correlated with those of proteins (**Figure 2B**). However, aging is the main risk of developing AD and one hallmark of aging is the loss of proteostasis. By decoupling mRNA and protein expression in monocytes, we could identify distinct subsets of genes that differentially adjust to the disease progression. A discordance between mRNA and protein abundance was recently described in neurodegenerative diseases ^16^, therefore assessing protein abundance by quantitative proteomic analyses in monocytes across the spectrum of AD could also reveal potential pathological protein drivers and foster new ways to investigate AD. Moreover, we previously described that the proportions of non-classical and intermediate monocytes increased through the spectrum of AD at the expense of classical monocytes ^7^. Single-cell methods have been extensively developed the last years and could help to further discriminate which monocyte population (e.g., non-classical vs classical monocytes) is targeted by AD-dependent mechanisms.

## Acknowledgements

This research was supported by a grant to B.L. from the Research Center on Aging at Sherbrooke. V.B. was supported by a fellowship from the Special Response Fund for Trainees (Ukraine) put in place by the three federal granting agencies i.e., the Canadian Institutes of Health Research (CIHR), the Natural Sciences and Engineering Research Council of Canada (NSERC) and the Social Sciences and Humanities Research Council of Canada (SSHRC). M.A.B. is supported by a Junior 1 fellowship from the Fonds de Recherche du Québec-Santé (FRQS).

## Conflict of interest

The authors have declared no competing interest.

